# Production of bioactive structural motifs from wheat arabinoxylan via colonic fermentation and enzymatic catalysis: evidence of interaction with toll-like receptors from *in vitro, in silico* and functional analysis

**DOI:** 10.1101/2024.09.13.612858

**Authors:** Caroline de A Guerreiro, Leandro AD Andrade, Layanne N Fraga, Tatiana M Marques, Samira BR Prado, Robert J Brummer, João Roberto O Nascimento, Victor Castro-Alves

## Abstract

It is well known that dietary fibers (DF) from plant-source foods can induce beneficial health effects through their physicochemical properties and utilization by the gut microbiota during fermentation, which is mainly explored with a focus on changes in the gut microbiota profile and the production of microbial-derived metabolites. Here, we characterized structural motifs (i.e., oligomers) produced during DF breakdown upon colonic fermentation and explored their interaction with toll-like receptors (TLRs) present on the surface of human intestinal and immune system cells. Firstly, a source of wheat arabinoxylan (WAX) was subjected to *in vitro* simulation of human colonic fermentation, followed by characterization and quantification of WAX structural motifs to explore their dynamics throughout fermentation. The identified structural motifs were further produced through enzymatic catalysis of WAX using carbohydrate-active enzymes and fractionated into six well-defined fractions of arabinoxylans and linear xylans. These fractions of structural motifs were then tested for interaction with TLR2 and TLR4 using a reporter cell assay. Results revealed structure-dependent effects, primarily with inhibition of TLR2 and activation of TLR4 depending on the degree of polymerization and branching of WAX structural motifs. The role of the fine structure of WAX structural motifs was confirmed by molecular docking, which revealed that minor structural changes substantially influence the interaction between structural motifs and TLRs. The results from *in vitro* and *in silico* studies also support the hypothesis that the direct effects of oligomers and polysaccharides on cell receptors are likely the result of complex interactions involving multiple cell surface receptors. Finally, in addition to highlighting that direct effects of structural motifs might play an important role in the overall effects of DF, this work suggests that enzymatic-tailoring design of DF can be a potential tool for producing functional ingredients with specific effects on human health.

## 1 Introduction

The intricate relationship between food ingredient structure, gut microbiota function and human health has evolved in ways that we are just beginning to understand. Despite questions that remain unclear, there is an increasing interest in exploring (and designing) bioactive food ingredients with improved health functionality. In this context, dietary fiber (DF) from plant-source foods accounts for the major DF fraction consumed by humans (An et al., 2022). Most of DF consist of complex cell wall-derived polysaccharides not hydrolyzed by human digestive enzymes. Among these, hemicelluloses, such as arabinoxylans (AX), are particularly important due to their presence in a variety of foods including wheat, rye, and barley (Zannini et al., 2022). These dietary sources contribute substantially to DF intake among different populations, being a focus of research in human nutrition and health. Chemically, AX consists of a backbone of β-(1→4)-linked xylose residues with side chains of arabinose linked to O-2 and/or O-3 positions of the xylose residues and, occasionally, other sugars. In some cases, ferulic acid residues and other neutral and acid sugars can be also linked to the arabinose side chains (Yao et al., 2023).

Despite the inability of human enzymes to break down the often-complex structure of DF including AX, their consumption is related with well-documented benefits due to physicochemical properties and effects associated with gut microbiota fermentation (Payling et al., 2020; Ye et al., 2022). The physicochemical effects are primarily associated with DF macrostructure, composed mostly of monosaccharides linked together, which chemically interact with water and organic compounds. This interaction results in a bulking effect, slowing absorption of soluble sugars and stimulating intestinal motility. Meanwhile, the effects associated with gut microbiota fermentation, occurring mainly at colon level, relates to the activity of carbohydrate-active enzymes (CAZy) produced by the gut microbiota to break down the DF structure, generating structural motifs that can serve as nutrients for their own metabolic processes or to cross-feed other microorganisms (Meng et al., 2022; Ye et al., 2022). This interaction between gut microbiota metabolism and microstructural aspects of AX, such as monosaccharide composition and linkage pattern, can lead to changes in both the abundance and diversity of the microbiome, as well as in changes in the levels of bioactive molecules including short-chain fatty acids (Rudjito et al., 2023) and tryptophan metabolites (Liikonen et al., 2024).

Beyond the production of microbiota-related metabolites, structural motifs from hemicellulose produced upon controlled acid hydrolysis or enzymatic catalysis of arabinans (Shi et al., 2020) can interact with the tool-like receptor (TLR)4 at the surface of intestinal epithelial cells (IEC) and immune system cells (Castro-Alves & Nascimento, 2021). This receptor, together with TLR2, has been associated with effects induced by DF (Beukema et al., 2020; Prado et al., 2020; Sahasrabudhe et al., 2018) and are among the first line of immune receptors expressed in human cells, being crucial for monitoring endogenous and exogenous components in the gut lumen (Duan et al., 2022). Thus, the interaction between TLRs and structural motifs produced either naturally via gut microbiota fermentation or commercially using physical processes or CAZy, have potential to boost host immune response, help to maintain intestinal immune homeostasis, and support therapy against intestinal immune disorders.

To explain a possible structure-function relationship between structural motifs from DF and cell receptors, which is highly desirable as a lead to effective industrial and translational applications, there is a need to both identify structural motifs produced upon colonic fermentation and to explore their synthesis and potential functionality. This knowledge will provide novel insights into the role of DF and bioactive oligosaccharides on health, thereby supporting the design of new functional ingredients. Our hypothesis is that structural motifs produced upon gut microbiota fermentation can contribute to the beneficial health effects of DF consumption by interacting with intestinal and immune system receptors. Here, we simulated human colonic fermentation of wheat arabinoxylan (WAX) to investigate the dynamics of structural motifs production throughout the fermentation process. Additionally, we explored enzymatic catalysis using specific CAZY to generate the same structures produced during colonic fermentation. To validate their potential functionality, structural motifs produced through enzymatic catalysis were isolated, purified, and tested for interaction with TLR2 and TLR4 using reporter cell assay. Finally, molecular docking was used to elucidate potential interaction mechanisms between bioactive WAX structural motifs and TLRs.

## 2 Materials and methods

### 2.1 Materials

Arabinoxylan from wheat flour (WAX, MW 56 KDa, Ara:Xyl = 38:62), α-L-arabinofuranosidase purified from *Aspergillus niger* (CAZy family: GH51, AFASE), endo-1,4-β-xylanase recombinant from *Neocallimastix patriciarum* (CAZy family: GH11, XYLNP), feruloyl esterase recombinant enzyme from rumen microorganism (CAZy family: CE1, FAERU), 3^2^-α-L-arabinofuranosyl-xylobiose (A^3^X), 2^3^-α-L-arabinofuranosyl-xylotriose (A^2^XX), ^23^-α-L-arabinofuranosyl-xylotetraose (XA^2^XX), 3^3^-α-L-arabinofuranosyl-xylotetraose (XA^3^XX), 2^3^,3^3^-di-α-L-arabinofuranosyl-xylotriose (A^2^,^3^XX), xylotriose (XTR), xylotetraose (XTE), xylopentaose (XPE) and xylohexaose (XHE) were purchased from NEOGEN/Megazyme (Auchincruive, Scotland). The human embryonic kidney (HEK 293) cell lines HEK-Blue hTLR2 and HEK-Blue hTLR4, Pam3CysSerLys4 (Pam3CSK4), lipopolysaccharide from *Escherichia coli* 0111:B4 (LPS), anti-hTLR2-IgA clone B4H2, anti-hTLR4-IgG clone W7C11 and HEK-Blue detection reagent were purchased from Invivogen (San Diego, USA). Cell culture consumables were purchased from Sartorius (Gottingen, Germany) and VWR International (Karlskoga, Sweden). Mass spectrometry-grade solvents were purchased from Fischer Scientific (Hampton, USA). Milled wheat bran (*Kruskakli*) containing 20% arabinoxylan (w/w) was kindly provided by Lantmännen Cerealia (Stockholm, Sweden). Other reagents were purchased from Sigma-Aldrich (Saint Louis, USA) unless stated otherwise.

### 2.2. Methods

The method section was divided in three subsections comprising simulation of colonic fermentation and enzymatic catalysis of WAX (*section 2.2.1*); identification, isolation and quantification of WAX structural motifs (*section 2.2.2*); and functional analysis of WAX structural motifs (*section 2.2.3*). An overview of the analysis workflow is illustrated in **Fig 1**.

**Fig 1.**
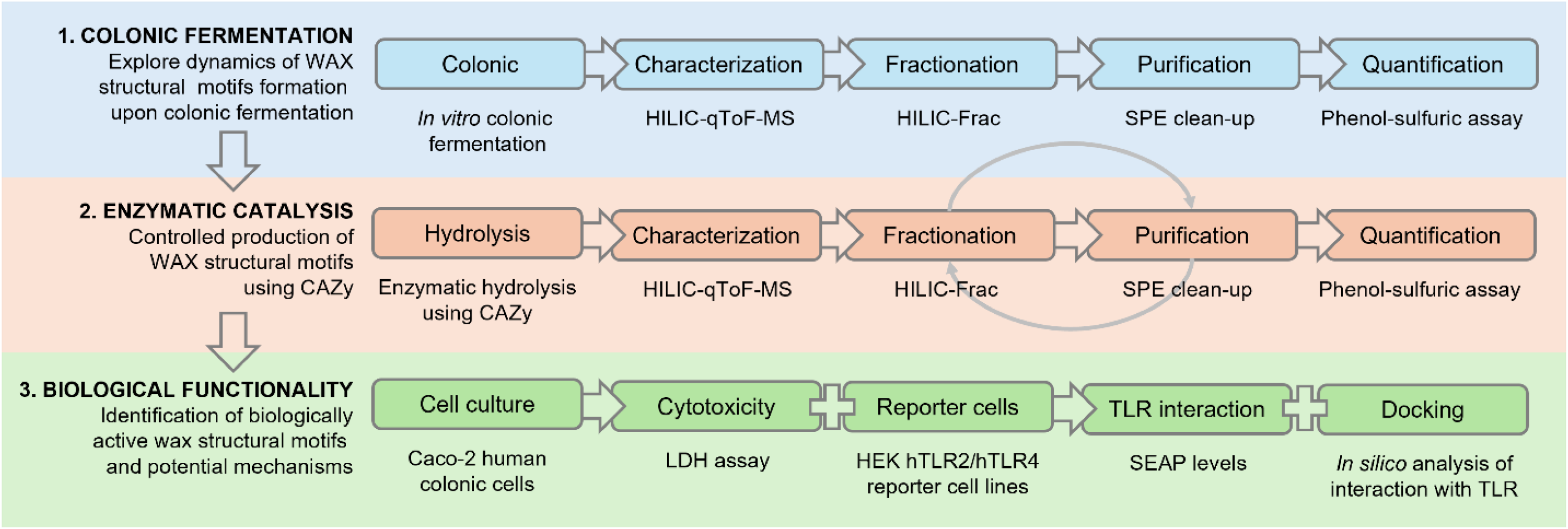
Analysis workflow. Structural motifs from wheat arabinoxylan (WAX) obtained upon colonic fermentation were characterized and further produced using carbohydrate-active enzymes (CAZy). Biological functionality of different WAX structural motifs was assessed using toll-like receptors (TLR)2 and TLR4 reporter cell lines after evaluation of potential cytotoxicity using Caco-2 human cells. Finally, molecular docking was applied to explore potential interaction mechanisms.

#### 2.2.1 Simulation of colonic fermentation and enzymatic catalysis of WAX

##### 2.2.1.1 In vitro simulation of human colonic fermentation

Three independent experiments each using the fecal inoculum of a healthy donor (two omnivorous females and one omnivorous male, ranging from 26 to 38 years-old) were performed to simulate human colonic fermentation. Experiments followed ethical guidelines and were approved by the Swedish Ethical Review Authority (#2022-01696-01). In each experiment a fecal sample was collected by the donor in a sterile plastic container and transported in an Oxoid AnaeroGen anaerobic pouch (Thermo Fisher). The fecal inoculum was prepared within 2 h of sample collection as described previously (Martín-Peláez et al., 2008). Briefly, a strainer stomacher bag with a filter was used to prepare the fecal inoculum by mixing 20 g of fecal sample (1:5, w/v) with 0.05% L-cysteine hydrochloride in 50 mM phosphate buffer. A 10 ml-aliquot of the inoculum was further mixed with 1 g of wheat bran (20% WAX, w/w; *Kruskakli*) and added to the fermentation flasks containing sterilized growth medium (1:10, v/v; pH 7) containing 0.2% peptone water, 0.2% yeast extract, 0.2% NaHCO_3_, 0.01% NaCl, 0.004% K2HPO_4_, 0.004% KH2PO_4_, 0.001% MgSO_4_, 0.004% CaCl_2_, 0.2% Tween 80, 0.005% hemin, 0.001% vitamin K1, 1.1% cysteine, 0.00002% resazurin, and 0.05% bile salts (Ox-bile). For each donor, one fermentation flask containing medium and inoculum without substrate (i.e., without wheat bran) was also prepared as a process fermentation blank (control). Aliquots (1 mL) collected from each fermentation flask after 0 (10 min), 2, 4 and 24 h were centrifuged (15000 x g, 10 min, 4 °C) and pH in the supernatants was recorded. Supernatants were stored at -80 °C for SCFA analysis and to explore dynamics of WAX structural motifs throughout the fermentation.

##### 2.2.1.2 Short-chain fatty acid (SCFA) analysis

Changes in the levels of acetate, propionate and butyrate throughout the fermentation were measured to confirm *in vitro* fermentation functionality. Samples were analyzed after derivatization with 3-nitrophenylhydrazine (3-NPH), similarly as described previously (Dei Cas et al., 2020). An ultra-high performance liquid chromatography coupled to time-of-flight high-resolution mass spectrometry (UHPLC-qToF-HRMS) was used for analysis. Briefly, a 40 µL-aliquot of sample was mixed with 480 µL of cold methanol/methyl-tert-butyl ether/isopropanol (MeOH/MTBE/IPA, 1.3:1:1) (Fu et al., 2022) containing 10 µg/mL acetic acid-d4, butyric acid-d8 and propionic acid-d2 as internal standards (IS). After sonication (5 min), the extract was incubated in an orbital shaker (1 h, 4 °C), filtered (0.45 µm), and a 25 µL-aliquot was subjected to derivatization with 3-NPH. Sample was further mixed with 50 mM 3-NPH (25 µL), 50 mM N-ethylcarbodiimide (EDC, 25 µL) and 7% v/v pyridine (25 µL). After incubation (1 h), 0.2% formic acid was added to stop the derivatization reaction and the sample was immediately analyzed using an Xevo G3 UHPLC-qToF-MS system (Waters Corporation, Milford, USA) equipped with an Acquity BEH C18 column (2.1 × 100 mm, 1.7 µm; Waters Corporation) connected to a pre-column (2.1 × 500 mm, 1.7 µm) and an 0.22 µm in-line filter. The injection volume was 3 µL. Mobile phase (MP)A containing 0.1 % formic acid in water and MPB containing acetonitrile were eluted at 0.4 mL/min starting with 10% MPB (0–2 min), 0–100% MPB (2–4 min), 100% MPB (4–6 min), followed by re-equilibration with 10% MPB for 4 min. The column and autosampler temperature were maintained at 50 °C and 10 °C, respectively. Accurate mass spectra were acquired (5 spectra/s) with a 50–1200 m/z range in negative ion mode. The source was programmed as follows: collision energy 0 V, capillary voltage 1.5 kV, sampling cone 40 V, desolvation temperature and gas flow at 550 °C and 16 L/min, and source temperature and cone gas flow at 150 °C and 0.8 L/min, respectively. Analysis was performed using the lock spray feature with leucine-enkephalin as reference at a frequency of 30 s throughout analysis. Data acquisition and pre-processing were performed using MassLynx v 4.1 (Waters Corporation) and MZmine 4.1.0 software (MZio; Bremen, Germany), respectively. Peak areas were normalized by the internal standards and results for acetate, butyrate and propionate were expressed as relative change (fold change) in relation to the first sampling point.

##### 2.2.1.3 Production of WAX structural motifs through enzymatic catalysis

Enzymatic catalysis of WAX was performed using α-L-arabinofuranosidase (AFASE), endo-1,4-β-xylanase (XYLNP) and feruloyl esterase (FAERU), used alone or in sequence to breakdown WAX aiming at releasing structural motifs. A total of five enzyme cocktails were tested (**Table S1**). Briefly, WAX was diluted (0.5 mg/mL, 10 mL) in an appropriate buffer and incubated with CAZy enzymes in accordance with manufacturer’s instructions (Megazyme). Samples were collected at 0.5 and 24 h after incubation with each enzyme. When more than one enzyme was used (i.e., cocktails #3, #4 and #5), the hydrolysate was dialyzed in between incubations using a 500 Da-MWCO biotech-grade cellulose dialysis tubing (Spectra-Por, Cole Parmer, Illinois, USA). The retentate was concentrated in a vacuum concentrator to further exchange buffer and adjust to optimal pH and temperature for enzyme activity. A total of 16 samples (five enzyme cocktails resulting in 8 incubation steps, each collected at two time points) were evaluated for the presence of WAX structural motifs. The selection of the best cocktails and incubation times for production of WAX structural motifs to proceed with functional analysis were based on the variety of structural motifs produced. In cases where different cocktails generated a similar profile, the enzyme cocktail able to produce the highest amount of hydrolysates was selected.

#### 2.2.2. Identification, isolation and quantification of WAX structural motifs

Samples obtained from *in vitro* simulation of human colonic fermentation of different donors and time points (n = 12) were analyzed for the presence of WAX structural motifs and further fractionated and purified for quantification. Samples collected from enzymatic catalysis (n = 16) were also explored for the presence of WAX structural motifs and, after selecting the best enzyme cocktails and time points, samples were fractionated and purified for quantification and to explore functionality using reporter cells.

##### 2.2.2.1 Target analysis of WAX structural motifs

The assessment of WAX structural motifs with degree of polymerization (DP) between DP3 to DP6 was performed by hydrophilic interaction liquid chromatography coupled to time-of-flight mass spectrometry (HILIC-qToF-MS), similarly as described previously (Amicucci et al., 2020; Juvonen et al., 2019). A 1290 Infinity ultra-high-performance system interfaced with a dual ESI source to a 6545 qToF-MS system (Agilent Technologies, Santa Clara, USA) and equipped with a porous graphitized carbon (PGC) Hypercarb column (100 × 2.1 mm, 3 µm) connected to a Hypercarb guard column (10 × 2.1 mm, 5 µm; Thermo Scientific, Waltham, USA) was used for analysis. Accurate mass spectra were acquired (2 spectra/s) with an m/z range of 70–1500 in positive ion mode. Mobile phase (MP)A 0.1% formic acid in water/acetonitrile (97:3, v/v) and MPB 0.1% formic acid (FA) in water/acetonitrile (10:90) were eluted at 0.3 mL/min using a stepwise gradient program starting with 20% MPB and increments of 10% MPB every minute until 6 min, followed by a re-equilibration step of 7 min. The injection volume was 5 µL. MassHunter Workstation Software (Agilent Technologies) was used for data acquisition. The identity of detected structural motifs in the hydrolysates was confirmed by matching mass-to-charge (m/z) of [M+Na]+ adducts and retention time (RT) to that of analytical standards of both AX residues and linear xylans (**Table 1**).

**Table 1.** Mass-to-charge ratio (m/z), retention time (RT, min) and response linearity of analytical standards used to identify and quantify structural motifs from wheat arabinoxylan (WAX). Analytical standards were analyzed in the same HILIC-qToF-MS analysis batch of the WAX hydrolysates obtained during colonic fermentation and enzymatic catalysis. Positive identification of peaks was determined by matching accurate m/z and RT with the following analytical standards. An additional standard containing a mixture of 2^3^- and 3^3^-α-L-arabinofuranosyl-xylotetraose (XA^2^XX/ XA^3^XX) was analysed to confirm the presence of two closely eluting oligomers with m/z 701.2112 at 4.99 and 5.16 min. The linearity refers to the calibration curve of each standard (4, 8, 12, 16, 20 and 40 µg/mL) using the phenol-sulfuric acid method (please refer to section 2.2.2.3).

| Analytical standard | m/z [M+Na]<sup>+</sup> |  | RT (min) | Linearity (r<sup>2</sup>) |
| --- | --- | --- | --- | --- |
|  | Theoretical | Experimental |  |  |
| 3<sup>2</sup>-α-L-Arabinofuranosyl-xyllobiose (A<sup>3</sup>X) | 437.1266 | 437.1267 | 3.32 | 0.9971 |
| 2<sup>3</sup>-α-L-Arabinofuranosyl-xylotriose (A<sup>2</sup>XX) | 569.1688 | 569.1688 | 4.15 | 0.9941 |
| 2<sup>3</sup>-α-L-Arabinofuranosyl-xylotetraose (XA<sup>2</sup>XX) | 701.2111 | 701.2112 | 4.99 | 0.9982 |
| 3<sup>3</sup>-α-L-Arabinofuranosyl-xylotetraose (XA<sup>3</sup>XX) | 701.2111 | 701.2112 | 5.16 | 0.9979 |
| 2<sup>3</sup>,3<sup>3</sup>-di-α-L-Arabinofuranosyl-xylotriose (A<sup>2,3</sup>XX) | 701.2111 | 701.2111 | 5.30 | 0.9984 |
| Xylotriose (XTR) | 437.1266 | 437.1269 | 3.56 | 0.9970 |
| Xylotetraose (XTE) | 569.1688 | 569.1692 | 4.48 | 0.9973 |
| Xylopentaose (XPE) | 701.2111 | 701.2115 | 5.48 | 0.9951 |
| Xylohexaose (XHE) | 833.2533 | 833.2535 | 5.72 | 0.9972 |

##### 2.2.2.2. Fractionation of WAX structural motifs

A HILIC fractionation system (Waters Corporation) controlled using the Empower Chromatography Data System software (Waters Corporation) with the same column and parameters applied for target analysis was used to collect fractions of WAX structural motifs into 96-well plates. Fractions were collected from 2.5 min to 6.5 minutes based on the chromatogram profile obtained for the structural motifs during HILIC-qToF-MS analysis. Each fraction was collected for 15 seconds throughout the collection time span, resulting in 16 fractions/sample that were further subjected to purification and quantification.

##### 2.2.2.3. Purification and quantification of WAX structural motifs

Purification of structural motifs fractions was performed using porous graphitic carbon (SPE-PGC) HyperSep Hypercarb SPE 96-well plates (Thermo Scientific) according to manufacturer’s instructions. A semi-automated robotic platform (Andrew+, Waters corporation) was designed using the OneLab online platform (https://onelab.andrewalliance.com/) for washing and conditioning the SPE-PGC resin, and for loading, washing and eluting the purified fractions into new 96-well sample collection plates. Purified fractions were dried to completion under vacuum centrifugation and reconstituted in 100 μL of LC-MS-grade water. A 50 µL-aliquot was used for quantification and characterization purposes, while the remaining was diluted in media for cell culture assay. The “.onp” file with the semi-automated protocol for purification of structural motifs is freely available to be uploaded to the OneLab online platform (a description of resources is available in the supplementary material).

Quantification was performed using the phenol-sulfuric method (Total Carbohydrate Assay Kit, Sigma-Aldrich). Briefly, 30 µL of the purified fractions and analytical standards at 4, 8, 12, 16, 20 and 40 µg/mL (described in **Table 1**) were transferred to a 1.5-mL polypropylene tube and mixed with reagent solution (5% w/v phenol) and concentrated sulfuric acid. Tubes were closed using a security clamp and incubated in an orbital shaker (20 min; 90 °C). After incubation, sample was maintained in an ice bath (5 min) and 100 µL was transferred to a 96-well plate for quantification. Absorbance was recorded at 490 nm using a FLUOstar Omega microplate reader (BMG Labtech, Ortenberg, Germany). Each fraction was analyzed in technical triplicate. Since the response of the phenol-sulfuric method is partially dependent on carbohydrate structure (Masuko et al., 2005), an aliquot of each fraction was re-analyzed by HILIC-qToF-MS to define the best analytical standard used for quantification. The degree of hydrolysis was calculated as the sum of WAX structural motifs obtained in relation to the amount of substrate (i.e., WAX) used during hydrolysis (%, w/w). The recovery of the SPE-PGC clean-up method (SPErec) was calculated as the sum of all structural motifs obtained after clean-up in relation to the sum of structural motifs before the SPE clean-up (%, w/w).

#### 2.2.3 Functional analysis of WAX structural motifs

After purification, fractions containing WAX structural motifs were tested for cytotoxicity at different concentrations in human intestinal epithelial cells and for interaction with human toll-like receptors (TLR)2 and TLR4 using reporter cell assay. Furthermore, molecular docking was performed to explain potential mechanisms associated with the interaction between bioactive structural motifs and TLRs.

##### 2.2.3.1 Cytotoxicity assay

The human intestinal epithelial cell line Caco-2 (ATCC HTB-37) was used to test potential cytotoxicity of the structural motifs. Briefly, cells were grown in DMEM supplemented with 10% heat-inactivated FBS, L-glutamine (2 mM), glucose (4.5 g/L), 100 U/mL penicillin and 100 μg/mL streptomycin (37 °C, 5% CO2). Viability was assessed with Trypan blue dye to ensure it was higher than 90% before cells were seeded in a 96-well plate (2.0 × 104 cells/well). The levels of lactate dehydrogenase (LDH) were measured in cell supernatant after 24-h incubation with the purified fractions containing the structural motifs (1, 10 and 25 µg/mL), cell culture media (non-treated cells) or 2% Triton X-100 (cell death control). Briefly, 50 µL of the supernatant was mixed with 50 µL of the reagent solution (Cytotoxicity Detection Kit, Sigma-Aldrich) and after 30 min absorbance was recorded at 490 nm. Results were expressed as the percentage of cell death in relation to cell death control. Each treatment was analyzed in triplicate in three independent experiments (n = 9).

##### 2.2.3.2 Reporter cell assay

A non-cytotoxic concentration was further selected to explore the interaction between the WAX structural motifs and TLR using HEK-Blue hTLR2 and HEK-Blue hTLR4 cell lines. These cells express an inducible reporter gene for SEAP (secreted embryonic alkaline phosphatase) upon activation of TLR2 and TLR4, allowing quantification of TLR response based on NF-κB activation followed by SEAP production, which is quantified using the Quanti-Blue reagent (InvivoGen). Briefly, cells were grown in DMEM supplemented with 10% heat-inactivated FBS, L-glutamine (2 mM), glucose (4.5 g/L), 100 U/mL penicillin, 100 μg/mL streptomycin, and 100 μg/mL normocin (37 °C, 5% CO2). After two passages, cells were maintained in a selective HEK-Blue selective medium. Viability was assessed with Trypan blue dye to ensure it was higher than 90% before they were seeded in a 96-well plate according to manufacturer’s protocol (2.5 -5.0 × 104 cells/well). For each reporter cell line, structural motifs were tested in three conditions:

a. “TLR activation”: cells were incubated with WAX structural motifs fractions or TLR agonists (24 h) and SEAP levels were measured in the supernatant. Agonists for TLR2 and TLR4 were Pam3CysSerLys4 (Pam3CSK4, 10 ng/mL) and LPS (10 ng/mL), respectively.
b. “Cell functionality”: cells were pre-treated with a blocking antibody for TLR2 or TLR4 one hour prior treatment with WAX structural motifs or TLR agonists (24 h). Structural motifs able to activate TLR (and TLR agonists) are expected to be inhibited by the pre-treatment with the blocking antibody—this condition was used to validate results from “TLR activation”. The inhibitors of TLR2 and TLR4 were anti-hTLR2-IgA clone B4H2 (10 µg/mL) and anti-hTLR4-IgG clone W7C11 (10 µg/mL), respectively.
c. “TLR inhibition”: reporter cells were incubated with a known agonist for TLR2 (10 ng/mL Pam3CSK4, Pam3CysSerLys4) or TLR4 (10 ng/mL LPS, lipopolysaccharide) one hour prior treatment with structural motifs and incubated for 24 h. Results were compared to reporter cells treated only with agonists. A decrease in agonist-mediated activation by a given fraction indicates the latter acts either by competitive inhibition or by blocking agonist activity.

After incubation in the conditions described above, 20 µL of cell supernatant was transferred to a 96-well plate containing 180 µL of QUANTI-Blue assay solution (InvivoGen) to measure SEAP levels. After 2 h, absorbance was recorded at 650 nm. Results from “TLR activation” and “Cell functionality” were expressed as the percentage of activation in relation to the TLR agonist (100% activation), while results from “TLR inhibition” were expressed as the relative inhibition in relation to cells incubated only with the TLR agonist (0%, no inhibition). Each treatment was analyzed in triplicate in four independent experiments (n = 12).

##### 2.2.3.3 Molecular docking

Molecular docking was performed to predict the strength of the molecular interactions that may occur between selected structural motifs and TLR2 and TLR4 using the SwissDock 2024 analysis tool (https://www.swissdock.ch/) (Bugnon et al., 2024; Grosdidier et al., 2011). The chemical structures of XTE (#10201852) and XPE (#10230811) were retrieved from PubChem (https://pubchem.ncbi.nlm.nih.gov/), while structures of A3X, A2XX, XA2XX, XA3XX and A2,3XX were modelled using the ChemDraw Professional v 16 software (PerkinElmer, Waltham, USA). The protein 3D structures of human TLR2 (#O60603) and TLR4 (#O00206) were retrieved from UniProtKB (https://www.uniprot.org/). Results were analyzed considering overall binding capacity to TLR2 and TLR4 scaffold. Interactions between -7.0 and -8.9 kcal/mol and lower than -9.0 kcal/mol were considered indicators of moderate and high binding capacity, respectively (Kitchen et al., 2004; Morris et al., 1998).

## 3. Results

### 3.1. Wheat arabinoxylan (WAX) structural motifs are produced upon colonic fermentation

Wheat bran flour with a known concentration of WAX was used as a substrate during *in vitro* simulation of human colonic fermentation as a proof-of-concept that WAX structural motifs are produced and can be quantified throughout the colonic fermentation process. The reduction in pH and increase in levels of SCFA acetate, propionate and butyrate were used as a proxy to confirm *in vitro* fermentation functionality. The fermentation using either WAX source as substrate or process blanks displayed a decrease in pH (**Fig S1A**) and increase in SCFA production throughout fermentation (**Fig S1B**). As expected, although the ratio of changes in pH and SCFA is different among donors, all samples from the same donor using WAX as substrate showed more pronounced changes compared to its process blank (i.e., no substrate added).

Following, the levels of selected WAX structural motifs were analyzed in the extracts obtained from *in vitro* colonic fermentation. After confirming the presence of structural motifs through HILIC-qToF-MS and matching mass-to-charge ratio and retention time to that of internal standards (**Table 1**), fermentation extracts were fractionated, purified and quantified. Although the limit of quantification (LOQ) of WAX structural motifs in our workflow ranged between 0.11 µg/mL and 27.76 µg/mL (**Table S2**), most of the structural motifs explored in this study were quantified in the extracts collected after 2 h of fermentation (**Table 2**). Results also revealed that the profile of structural motifs changes over time throughout fermentation and exhibits different dynamics among participants. The ratio between structural motifs containing side arabinose residues and linear xylans (AX:X ratio) was calculated as an indicative of changes in dynamics within and between donors, mostly showing an increase in AX:X ratio when comparing 4 h and 24 h of incubation. These results connected to the below-LOQ (or non-detection) levels of structural motifs in process blanks (**Table S3**) confirm that the detected structural motifs were mainly a result of substrate (i.e., WAX) fermentation by the gut microbiota.

**Table 2.** Dynamics of WAX structural motifs throughout *in vitro* simulation of human colonic fermentation. Structural motifs were quantified after purification, annotation, fractionation and concentration. Results are expressed as µg/ 100 mL of the structural motifs in the fermentation extract (i.e., total amount/fermentation flask) and, in parentheses, as the relative amount (%, v/v) in relation to the amount of WAX used during fermentation (approx. 2 mg WAX/fermentation flask). Fermentation process blanks (i.e., without WAX source) were also explored for the presence of structural motifs (**Table S3**). LOQ: value below the limit of quantification (**Table S2**); -: not detected; AX:X ratio: ratio between levels of structural motifs containing arabinose units and linear xylans.

| Oligomer fractions | Donor 1 |  |  |  | Donor 2 |  |  |  | Donor 3 |  |  |  |
| --- | --- | --- | --- | --- | --- | --- | --- | --- | --- | --- | --- | --- |
|  | 0 h | 2 h | 4 h | 24 h | 0 h | 2 h | 4 h | 24 h | 0 h | 2 h | 4 h | 24 h |
| A<sup>3</sup>X | LOQ | 18.1 (0.9) | 26.7 (1.3) | LOQ | LOQ | 16.0 (0.8) | - | - | - | 28.8 (1.4) | 30.9 (1.5) | LOQ |
| A<sup>2</sup>XX | LOQ | 1.3 (0.1) | 0.5 (<0.1) | - | - | - | 34.0 (1.7) | LOQ | 19.6 (1.0) | 54.2 (2.7) | 41.8 (2.1) | LOQ |
| XA<sup>3</sup>XX | 77.5 (3.9) | 66.3 (3.3) | 96.1 (4.8) | 166.6 (8.3) | 40.1 (2.0) | 62.4 (3.1) | 110.3 (5.5) | 134.7 (6.7) | 66.6 (3.3) | 85.2 (4.3) | 122.8 (6.1) | 96.1 (4.8) |
| A<sup>2,3</sup>XX | 58.3 (2.9) | 67.3 (3.4) | 106.5 (5.3) | 104.4 (5.2) | 28.3 (1.4) | 60.9 (3.0) | - | - | 62.1 (3.1) | 71.5 (3.6) | 120.4 (6.0) | 121.3 (6.0) |
| XTE | LOQ | 42.0 (2.1) | 66.7 (3.3) | 76.4 (3.7) | - | 41.2 (2.1) | 149.3 (7.5) | 69.6 (3.5) | - | 59.5 (3.0) | 88.7 (4.4) | 51.4 (2.6) |
| XPE | - | 35.2 (1.8) | 49.2 (2.5) | 42.3 (2.1) | - | 40.7 (2.0) | 90.8 (4.5) | 60.1 (3.0) | - | 47.8 (2.4) | 39.5 (2.0) | 34.9 (1.8) |
| AX:X ratio | - | 2.0 | 2.0 | 2.3 | - | 1.7 | 0.6 | 1.0 | - | 2.2 | 2.5 | 2.6 |
| Total | 135.8<br>(6.2) | 230.2<br>(11.6) | 345.6<br>(17.2) | 389.7<br>(18.6) | 68.4<br>(2.7) | 221<br>(11.0) | 384.4<br>(19.2) | 264.4<br>(13.0) | 148.3<br>(7.4) | 347<br>(17.4) | 444.1<br>(22.1) | 300.7<br>(14.0) |

### 3.2 Production of WAX structural motifs through enzymatic catalysis

Enzymatic catalysis of WAX using carbohydrate-active enzymes (CAZy) was applied to generate the same structural motifs detected upon colonic fermentation of WAX to further test their biological functionality. Three CAZy were used alone or in combination to breakdown WAX structure. We did not detect any of the selected structural motifs when WAX was incubated with α-L-arabinofuranosidase alone (AFASE, cocktail #1). On the other hand, incubation with endo-1,4-β-xylanase alone (XYLNP, cocktail #2) yielded four main groups of structural motifs after 24 h of incubation (**Fig 2A**). For cocktail #3 (AFASE + XYLNP), we did not detect structural motifs during incubation with AFASE; however, sequential incubation with XYLNP for 30 min resulted in the formation of distinct isomeric structures to those detected during incubation with XYLNP alone (**Fig 2B**). Incubation with AFASE followed by XYLNP for 24 h appears to lead to extensive hydrolysis of WAX as we did not detect structural motifs. We were not able to quantify structural motifs when FAERU was added before AFASE (cocktail #4). Furthermore, when FAERU was added before XYLNP (cocktail #5), results were similar to when XYLNP was used alone. These findings suggest that WAX used in this study has a low degree of esterification. Based on these results, the hydrolysates obtained from XYLNP (cocktail #2) after 24 h and from sequential incubation with AFASE for 24 h and XYLNP for 30 min (cocktail #3) were selected as the enzyme cocktails and time points to proceed with generation WAX structural motifs.

**Fig 2.**
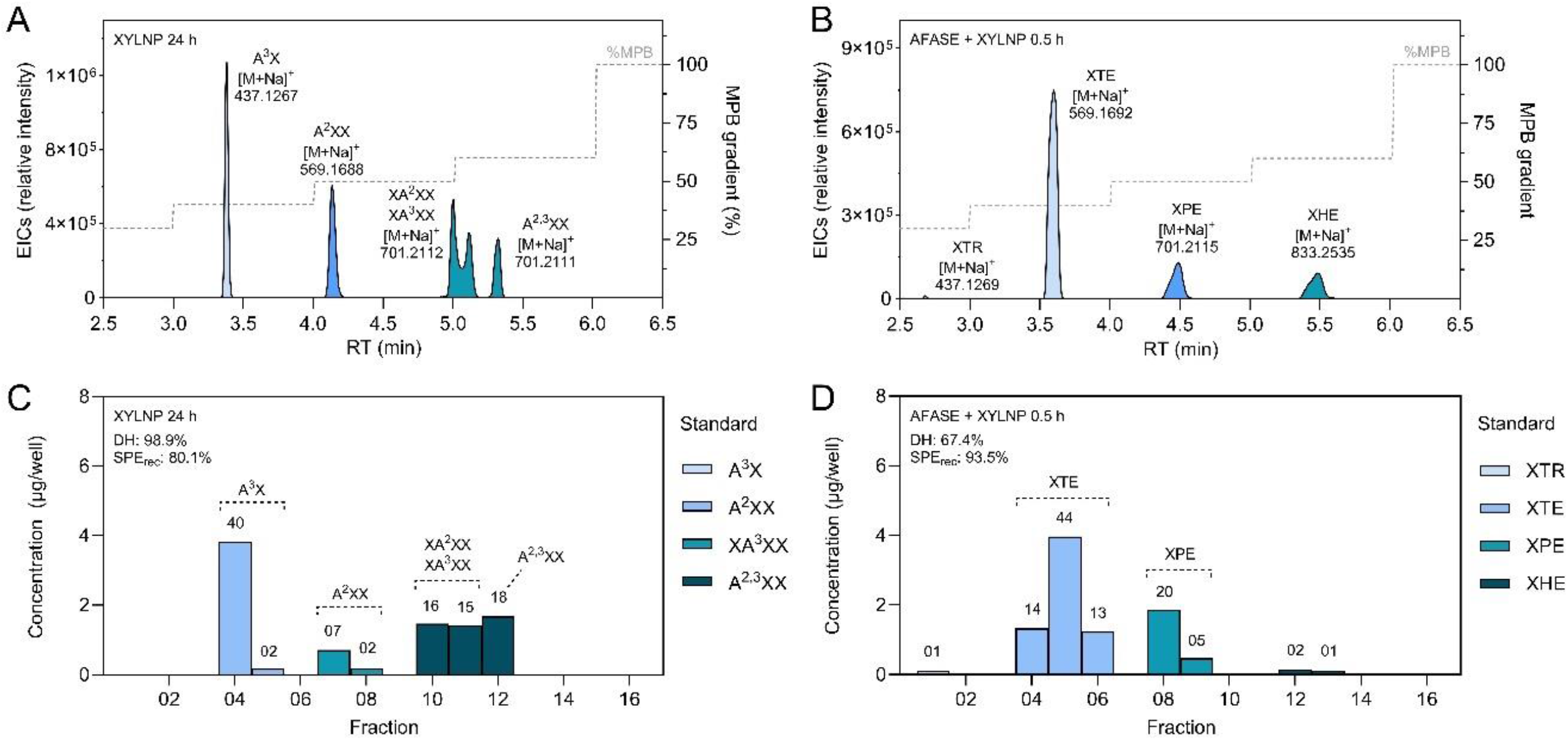
Structural motifs of wheat arabinoxylan (WAX) produced upon enzymatic catalysis. **(A)** Incubation with endo-1,4-β-xylanase (XYLNP) and with (**B**) α-L-arabinofuranosidase (AFASE) followed by XYLNP were selected to produce structurally diverse WAX structural motifs. (**C**) A total of six groups of structural motifs were selected for functional analysis, being four (A^3^X, A^2^XX, XA^2^XX/XA^3^XX and A^2,3^XX) obtained after incubation with XYLNP alone, and (**D**) two (XTE and XPE) obtained after sequential incubation with AFASE and XYLNP. The names above chromatograms and bars refer to the identity of the structural motif after confirmation of accurate m/z and retention time using analytical standards. The numbers above bars represent the relative amount of oligomer (%, w/w) in the fraction in relation to the amount of oligomer in all fractions. Different colours represent the analytical standard used for identification and quantification. EIC: extracted ion chromatogram; %MPB: mobile phase gradient used during the HILIC-qToF-MS analysis. DH: degree of hydrolysis (i.e., relative amount of oligomers obtained in relation to the amount of WAX used for hydrolysis); SPErec: PGC-SPE clean-up recovery (i.e., relative amount of oligomers quantified after purification in relation to the amount of oligomers obtained during hydrolysis).

As shown in **Fig 2C**, four groups of WAX structural motifs were collected when incubating WAX with XYLNP alone. As we were unable to clearly separate 23- and 33-α-L-Arabinofuranosyl-xylotetraose (XA2XX and XA3XX), these structural motifs were grouped into a single fraction for further analysis. Comparison between the quantification of fractions before and after the purification process indicates good recovery (SPErec: 80.1%). Furthermore, in addition to four fractions of arabinoxylans, we obtained two fractions of linear xylans, named xylotetraose (XTE) and xylopentaose (XPE), when incubating WAX sequentially with AFASE and XYLNP (**Fig 2D**). Xylotriose (XTR) and xylohexaose (XHE) were also detected, but functional analysis was performed only with fractions that could provide at least 5% of the selected WAX structural motifs due to the relatively low recovery associated with the low degree of hydrolysis (DH: 37.4%). Thus, we proceeded with functional analysis of six fractions of structural motifs from WAX displaying a varied degree of polymerization and representing a variety of branched and linear structures, as follows: (i) 3^2^-α-L-Arabinofuranosyl-xylobiose (A^3^X, and possibly A^2^X), (ii) 2^3^-α-L-Arabinofuranosyl-xylotriose (A^2^XX), (iii) a mixture of XA^2^XX and XA^3^XX, (iv) 2^3^,3^3^-di-α-L-Arabinofuranosyl-xylotriose (XA^2,3^XX), (v) XTE and (vi) XPE.

### 3.3 WAX structural motifs interact with human toll-like receptors in a structure-dependent manner

After isolating six fractions of structural motifs of WAX, we tested their interaction with TLR2 and TLR4 using HEK293-Blue hTLR2 and hTLR4 reporter cell lines. A preliminary cytotoxicity assay was performed using the human intestinal cell line (CaCo-2), to ensure observed results were associated with a direct biological function rather than an indirect effect caused by cytotoxicity. As shown in **Table 3**, all fractions of WAX structural motifs were not cytotoxic at the tested concentrations (1, 10 and 25 µg/mL). Thus, we proceeded with tests with reporter cells using 25 µg/mL.

**Table 3.** Effects of structural motifs from wheat arabinoxylan (WAX) on cytotoxicity. intestinal epithelial Caco-2 cells were treated with WAX or oligomer fractions (1, 10 and 25 µg/mL) and levels of lactate dehydrogenase (LDH) released, which are indicative of cytotoxicity, were compared to that of non-treated cells (control). Results are expressed in % of LDH levels in relation to 0.1% Triton X-100, used as a cell death control. No significant differences were observed between treatments and negative control (ANOVA, Dunnett’s post hoc test; p < 0.05). Results represent the mean ± SD of at least three independent experiments performed in triplicate (n = 9).

| Group | LDH release in comparison to cells treated with cell death control |  |  |
| --- | --- | --- | --- |
| Negative control (no treatment) | 11.5 ± 3.6 | | |
| Treatment | Concentration (µg/mL) |  |  |
|  | 1 | 10 | 25 |
| WAX | 9.8 ± 4.2 | 9.7 ± 2.1 | 9.5 ± 3.9 |
| A<sup>3</sup>X | 6.1 ± 1.5 | 6.9 ± 2.8 | 5.6 ± 2.8 |
| A<sup>2</sup>XX | 5.6 ± 3.1 | 4.2 ± 4.0 | 6.4 ± 3.3 |
| XA<sup>2</sup>XX/XA<sup>3</sup>XX | 5.9 ± 4.4 | 5.8 ± 3.9 | 7.1 ± 4.2 |
| XA<sup>2,3</sup>XX | 11.8 ± 5.2 | 7.6 ± 1.7 | 8.5 ± 3.1 |
| XTE | 10.6 ± 3.5 | 13.9 ± 6.8 | 12.9 ± 5.2 |
| XPE | 9.0 ± 3.8 | 14.5 ± 6.4 | 15.0 ± 4.5 |

As shown in **Fig 3A**, WAX structural motifs were not able to activate TLR2 at the tested conditions, while pre-incubation of cells with TLR2 blocking antibody inhibited activation by Pam3CSK4, a synthetic triacylated lipopeptide used as TLR2 agonist, confirming cell reporter assay functionality (**Fig 3B**). On the other hand, structural motifs significantly reduced TLR2 activation by Pam3CSK4 (**Fig 3C**). Among arabinoxylans (A^3^X, A^2^XX, XA^2^XX/XA^3^XX and XA^2,3^XX), the higher the degree of branching and polymerization, the higher the degree of inhibition; while linear xylans, XTE and XPE, had reduced and no inhibitory activity, respectively, at the tested concentrations.

**Fig 3.**
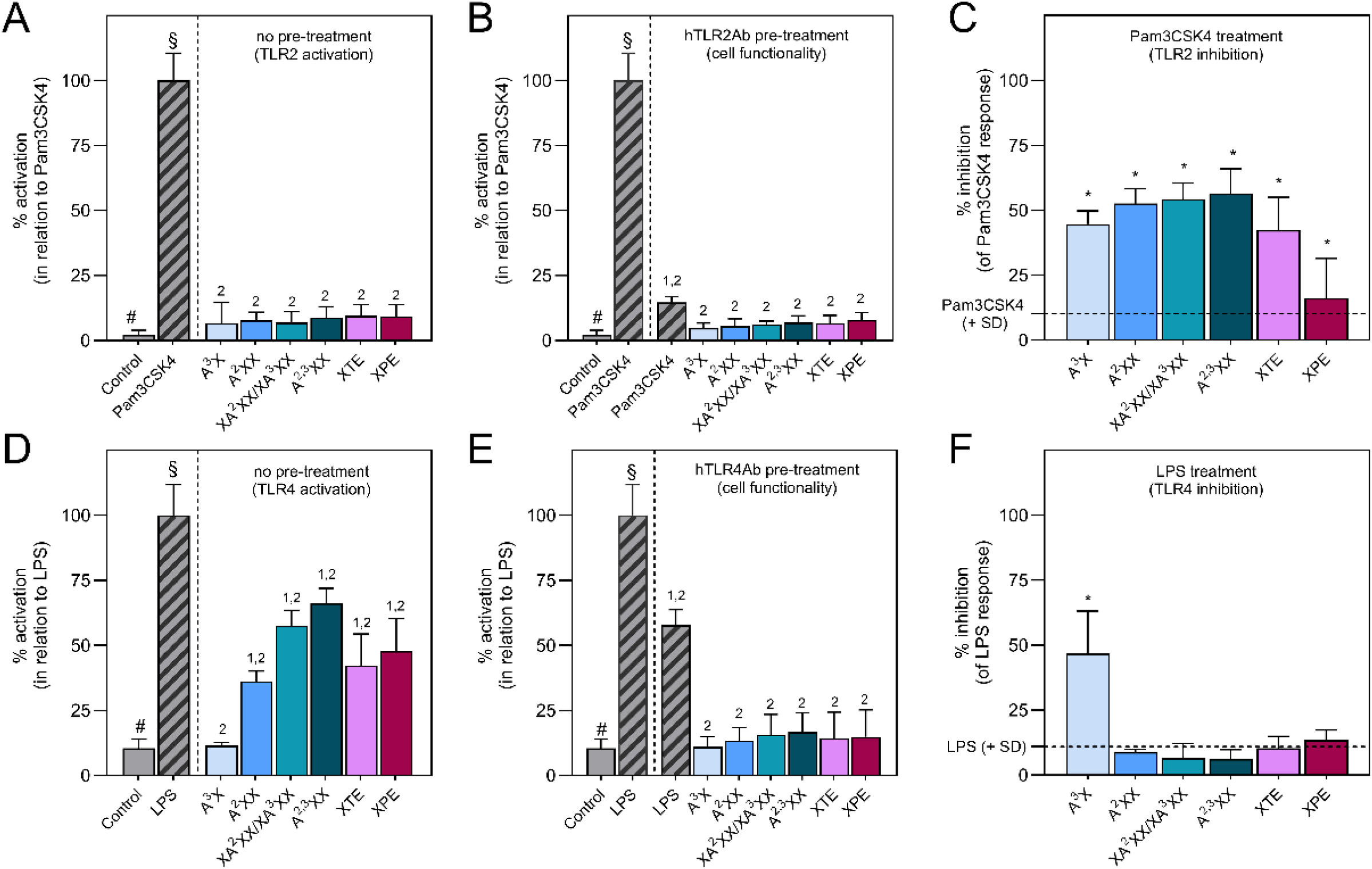
Effects of structural motifs from arabinoxylan (WAX) on toll-like receptors (TLR)2 and TLR4. NF-κB–SEAP reporter HEK293 cells expressing human TLR2 or TLR4 was incubated with WAX oligomers, TLR2 agonist (Pam3CSK4, Pam3CysSerLys4) or TLR4 agonist (LPS, lipopolysaccharide from *E. coli*) for 24 h to explore (**A**) TLR2 and (**D**) TLR4 activation. Samples were also incubated with cells pretreated (**B**) human blocking antibody for TLR2 (hTLR2Ab) or (**E**) TLR4 (hTLR4Ab) for 1 h to explore reporter cell functionality. Additionally, WAX oligomers were also (**C**) incubated one hour after Pam3CSK4 or (**F**) LPS to explore potential inhibitory effects of WAX oligomers compared to cells treated with only the agonist. To explore TLR activation and cell reporter functionality (**A, B, D** and **E**), results were calculated in relation to cells treated with only the agonist (100% activation). To explore TLR inhibition (**C** and **F**), results were expressed as the relative reduction in TLR activation compared to the agonist (0% inhibition); dashed line represents the SD interval obtained for the agonist. ^1^: significant difference in relation to untreated cells (control, #; ANOVA; Dunnett’s post hoc test; p < 0.05); ^2^: significant difference in relation to cells treated with Pam3CSK4 or LPS (§; ANOVA; Dunnett’s post hoc test; p < 0.05). *: significant inhibition in comparison to the agonist (ANOVA; Dunnett’s post hoc test; p < 0.05). Results represent the mean ± SD of at least four independent experiments performed in triplicate (n = 12).

Conversely to effects on TLR2, most of the WAX structural motifs significantly activated TLR4 (**Fig 3D**), and pre-incubation with TLR4 blocking antibody inhibited structural motif-induced activation, thereby confirming cell reporter functionality (**Fig 3E**). Among arabinoxylans (A^3^X, A^2^XX, XA^2^XX/XA^3^XX and XA^2,3^XX), the higher the degree of branching and polymerization, the higher the degree of TLR4 activation. As shown in **Fig 3F**, when structural motifs were incubated before LPS, a TLR4 agonist, there was no significant inhibitory activity, except for A^3^X, which inhibited LPS-induced activation by 46.7 ± 16.5%.

Finally, molecular docking was applied to elucidate potential interaction mechanisms between bioactive WAX structural motifs and TLRs. As shown in **Table 4**, the structural motifs exhibited moderate (between -7.0 and -8.9 kcal/mol) to high (< -9.0 kcal/mol) binding capacities when interacting with the entire TLR2 and TLR4 structures. The potential binding affinity of WAX structural motifs appears to increase with the degree of polymerization (DP), ranging from -7.6 to -9.1 kcal/mol. Notably, even though WAX structural motifs analyzed in this study share similar structural features, docking analysis reveals they have better affinities for different regions of the TLR scaffold, as highlighted by the different docking of A3X, A^2^XX, XPE, and XTE with TLR2 and TLR4 in **Fig 4**. A detailed representation of docking results including all WAX structural motifs analyzed in this study is available in **Fig S2**. This latter data also supports the hypothesis that slight changes in the structure of molecular motifs can lead to differences in their interaction with TLR2 and TLR4.

**Table 4.** Biding capacity of structural motifs from arabinoxylan (WAX) with the tool like receptor (TLR)2 and TLR4. Results represent the lowest binding energy (i.e., better binding affinity) to TLR2 and TLR4. XA^2^XX and XA^3^XX belong to the same structural motif fraction but were analyzed separately to explore whether differences in the spatial conformation of structural motifs impact their binding capacity.

| Structural motif | $\Delta G$ Binding/Docking<br>(kcal/mol) | |
| --- | --- | --- |
|  | TLR2 | TLR4 |
| A<sup>3</sup>X | -7.60 | -8.18 |
| A<sup>2</sup>XX | -7.74 | -9.12 |
| XA<sup>2</sup>XX | -8.43 | -8.64 |
| XA<sup>3</sup>XX | -9.08 | -9.27 |
| A<sup>2,3</sup>XX | -8.36 | -8.66 |
| XTE | -7.77 | -8.39 |
| XPE | -8.46 | -8.62 |

**Fig 4.**
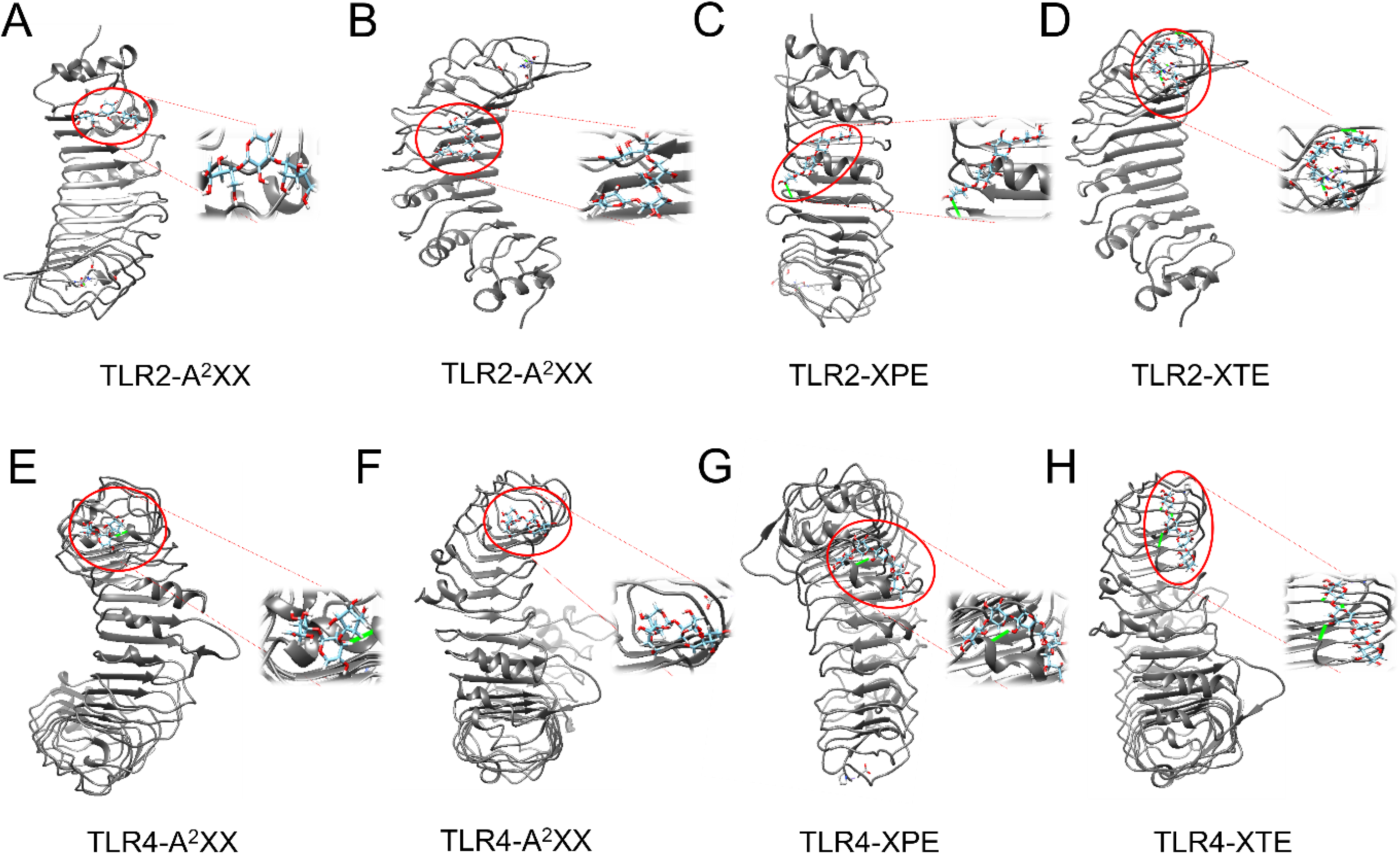
Molecular docking of structural motifs from wheat arabinoxylan (WAX) with toll-like receptors (TLR)2 and TLR4. Representative image of the highest score position (i.e., higher binding affinity) between A^3^X, A^2^XX, XPE and XTE in the molecular docking model with (**A-D**) TLR2 and (**E-H**) TLR4. Representative images of all structural motifs are shown in **Fig S2**.

## 4. Discussion

In this study, we applied a bioanalytical platform to characterize, generate and explore the biological effects of structural motifs produced upon gut microbiota fermentation of wheat arabinoxylan (WAX). We revealed that WAX structural motifs produced upon colonic fermentation display structure-dependent effects in the human intestinal and immune receptors TLR2 and TLR4. On a broader perspective, by combining analytical approaches to characterize and purify WAX structural motifs, with simulation of human colonic fermentation, enzymatic catalysis, and *in vitro* and *in silico* functional studies, we established a pipeline to explore (and validate) potential mechanisms of bioactive structural motifs. Previous studies by our group and others have shown biological effects of commercially available oligosaccharides, such as xylo- and arabino-oligosaccharides (Batsalova et al., 2022; Castro-Alves & Nascimento, 2021), while others have been exploring biological effects of complex oligosaccharide mixtures on human intestinal and immune response (Fernández-Lainez et al., 2022; Yasuma et al., 2021). However, to the best of our knowledge, this study presents a pioneering approach combining characterization of structural motifs produced upon colonic fermentation with their production and isolation via enzymatic catalysis, and further assessment of their biological potential combining simulation of human colonic fermentation and *in silico* approaches.

Firstly, simulation of colonic fermentation was performed as a proof-of-concept that structural motifs can be produced from a WAX source (in this case wheat bran) upon human gut microbiota fermentation. We found WAX structural motifs up to 24 h fermentation in the fermentation extracts of all donors and observed a distinct profile of structural motifs among donors. These findings were somewhat expected, given the individual diversity of the gut microbiota profile and functioning, which play a crucial role in shaping intraindividual fermentation kinetics. In another study exploring arabinoxylans, Pollet et al. (2012) focused on exploring the prebiotic potential of arabinoxylans from different sources and revealed that a significant proportion of arabinoxylan is still available after 48 h of colonic fermentation, especially when wheat bran was used as the substrate. Such findings also support our hypothesis that structural motifs from WAX can be available in the colon and thus contribute to the observed beneficial health effects in humans associated with WAX consumption (Boll et al., 2016; François et al., 2014).

While our study explored the structure-function relationship of structural motifs, related research has been focusing on the metabolization of DF by identifying specific microorganisms and the association between gut microbiota profile and CAZy enzymes, both at gene and functional levels (Fonseca et al., 2020; Ravn et al., 2021). Particularly for WAX, it has been shown that bacteria from the *Prevotella* genus appear to have an important role in the production of CAZy involved in WAX metabolization, such as α-L-arabinofuranosidase (AFASE) (Aakko et al., 2020; Chung et al., 2020), while feruloyl esterase and most of xylanase-containing isolates from healthy human feces appear to belong to the *Bacteroides* genus (Pereira et al., 2021; Zhang et al., 2022). Although our focus in simulating human colonic fermentation was to explore the possibility of recovering and quantifying WAX structural motifs, regardless of individual gut microbiota profiles and CAZy enzyme activity, knowledge of these latter is crucial for understanding individual responses to the utilization of DF by the gut microbiota and production of bioactive structural motifs.

The structures of arabinoxylans are relatively complex and source-dependent, but mostly composed of a backbone of β-(1→4)-linked xylose residues, which is occasionally branched with xylose at either O-3 position or both at O-2 and O-3 positions of the xylose residues (Rudjito et al., 2023; Wang et al., 2020). Thus, we selected an analytical approach able to characterize mainly small and non-esterified arabinoxylans, as well as linear xylans. We observed a relatively low degree of hydrolysis when using AFASE, either alone or in combination with XYL, which suggests there is greater potential for generating bioactive structural motifs from arabinoxylan by improving catalysis processes. Indeed, from a translational perspective, in addition to exploring the potential biological effects of structural motifs, there is a need to optimize enzymatic catalysis processes (Bhattacharya et al., 2020). Here, commercially available enzymes were used to break down WAX structure at defined sites, successfully generating the same structural motifs that were detected upon simulation of colonic fermentation. However, scaling-up this process using purified CAZy would most likely generate costs that at the current state are not feasible for commercial applications. One solution to circumvent this challenge would be to use either non-enzymatic treatment (Yue et al., 2022) or precision fermentation, this later by selecting or modifying microbial strains to express specific CAZy capable of producing targeted bioactive oligosaccharides on a large scale. Another strategy would be to explore associations between physiological parameters related to digestion (e.g., CAZY activity in feces, gut transit time) that could be used to predict the formation of specific structural motifs upon ingestion of a given DF. In this latter strategy, the gut microbiota machinery of the host would be responsible for the generation of the structural motifs. From a biotechnological perspective, studies have been exploring potential cell factories that can be used to tailor the structure of DF and other food components, as recently reviewed (Yang et al., 2023). For arabinoxylans, recent research on enzymatic catalysis of different sources has identified promising microbial strains and enzymes that can be used to modify WAX structure, offering the potential to generate a diverse range of oligosaccharides (Rudjito et al., 2023).

Our findings showed that WAX structural motifs produced upon gut microbiota fermentation interact with TLR2 and TLR4, providing support for strategies aiming at tailoring DF structure to produce functional DF with beneficial (and desired) health effects. We found that most of the WAX structural motifs appear to inhibit TLR2 and activate TLR4. Nevertheless, there is still a need to understand the implications of these interactions on human health. Previous studies revealed that inhibition of TLR2 response by DF has been associated with protective effects against ileitis (Sahasrabudhe et al., 2018), while activation of TLR4 and the NF-κB downstream pathway has been associated with protective effects on intestinal barrier function (Baggio et al., 2022; Vogt et al., 2016). In line with these findings, a previous study involving over sixty volunteers showed that consumption of WAX was associated with improvements in gastrointestinal health parameters (François et al., 2012) and glucose tolerance (Boll et al., 2016). While the authors suggested that positive effects of WAX were linked to changes in gut microbiota and increased production of SCFA, our findings, along with studies showing how direct modulation of TLR by DF influences gastrointestinal health, suggest that mechanisms beyond changes in microbiota and SCFA levels may also be in place. Thus, it is crucial to explore the direct implications of structural motifs produced upon colonic fermentation to fully understand their contribution to the health benefits associated with DF consumption.

While highlighting the potential of WAX structural motifs to induce direct effects, our study also provides a bioanalytical approach to explore additional mechanisms. For instance, exploring the potential bioavailability of oligosaccharides and polysaccharides has been analytically challenging. It is well-known that DF can be internalized by immune system cells. Moreover, in the last decade, studies have been trying to explore transport mechanisms of polysaccharides through the gastrointestinal tract (Zuo et al., 2016) and distribution of DF—or their resulting structural motifs—in different tissues (Shao et al., 2022). In this later study, authors reported the tissue distribution of an inulin-type fructan upon oral administration in mice; however, analyses were performed after derivatization of the fructan with a fluorescent tag, which may influence the internalization and distribution of the derivatized molecule. By allowing isolation and quantification of specific structural motifs, our approach holds promise for advancing studies that aim to explore absorption and distribution of oligosaccharides without the need for prior chemical modification. Furthermore, the approach presented here can be used to recover oligosaccharides from biological matrices or from *in vitro* and *ex vivo* permeability studies, thereby allowing transmembrane transport studies and analysis of bioactive oligosaccharides *in situ*.

Finally, in addition to using reporter cells to explore the interaction between WAX structural motifs and TLR2 and TLR4, we also applied molecular docking to explore potential interaction mechanisms between WAX structural motifs and TLRs. An extensive retrospective analysis exploring the structure of DF that interacts with TLR4 suggests that oligo- and polysaccharides could simultaneously interact with multiple cell surface receptors (Li et al., 2022). More recently, authors used *in vitro* and *in silico* analysis to confirm this hypothesis by exploring the interaction of glucomannans with TLR4 and other surface receptors from immune system cells, for instance, CD14 and mannose receptor (MR), suggesting that TLR4-mediated responses are rather a multi-receptor synergistic immune response (Li et al., 2023). Our results from *in vitro* and *in silico* analysis confirm that structural motifs from WAX also interact with different receptors at the cell surface, for instance, TLR2 and TLR4. The observation of moderate to strong binding capacity of WAX structural motifs to different regions of the TLR2 and TLR4 scaffold supports the hypothesis that DF—and their structural motifs—can mediate their effects through interactions with multiple receptors at the cell surface. These interactions may involve activation of binding pockets and changes in receptors’ functional state by engaging with the protein scaffold. This apparent promiscuity of oligo- and polysaccharides may also explain the complexity of defining the structure-function relationship of the often-complex DF structures.

## 5 Conclusion

Using a bioanalytical platform, we identified structural motifs produced during human colonic fermentation of WAX and explored their interaction with surface receptors of intestinal and immune system cells. Interestingly, variations in the degree of polymerization and structure of isomeric structural motifs significantly impact their interaction with TLR2 and TLR4. The higher the degree of branching and polymerization of arabinoxylan, the more pronounced TLR2 inhibition and TLR4 activation, while linear xylans had reduced and no TLR2 inhibitory activity and less pronounced effects on TLR4 activation compared to arabinoxylans. These findings are in line with other recent studies and suggest that the health benefits of DF extend beyond physicochemical and prebiotic effects, highlighting the importance of considering direct effects at cellular level throughout the gastrointestinal tract to fully explain the beneficial effects of DF on human health. Considering the individual nature of the digestion process and gut microbiota functioning, it will be not surprising to verify a high variability among individuals in the capacity to produce bioactive structural motifs. Thus, information on the physiological aspects associated with the digestion process that drive the formation of bioactive structural oligomers needs to be explored. Furthermore, studies on the definition of the structure-function relationship between structural motifs and human cell receptors, as well as downstream biological responses of such interaction, can support the tailored production of DF with specific and desired effects on human health.

## CRediT authorship contribution statement

**Caroline Guerreiro:** Methodology, Investigation, Writing - Review & Editing. **Leandro Andrade:** Methodology, Formal Analysis, Investigation. **Layanne Fraga:** Methodology, Software, Formal Analysis. **Tatiana Marques:** Methodology, Investigation, Resources, Writing - Review & Editing. **Samira Prado:** Conceptualization, Investigation, Writing - Review & Editing, Visualization. **Robert Brummer:** Conceptualization, Resources, Writing - Review & Editing. **João Roberto Nascimento:** Conceptualization, Writing - Review & Editing, Supervision. **Victor Castro-Alves:** Conceptualization, Formal Analysis, Data Curation, Writing - Original Draft, Supervision, Project administration, Funding acquisition

## Data availability

The data that support the findings of this study are available from the corresponding author, VCA, upon request.

## Acknowledgments

We acknowledge Daniel Duberg and Eleftheria Gallou for their technical support. We also thank the Research Internship Abroad program of the São Paulo Research Foundation (BEPE-FAPESP, #2022/08480-0) for providing financial support to Leandro Andrade. This work was funded by a Starting Grant within Natural and Engineering Sciences of the Swedish Research Council (Project grant 2021- 04937) and the Lantmännen Research Foundation (Project grant 2022H004).

