## Supporting information for "Production of bioactive structural motifs from wheat arabinoxylan via colonic fermentation and enzymatic catalysis: evidence of interaction with toll-like receptors from *in vitro, in silico* and functional analysis"

### SUPPLEMENTARY MATERIAL

**CONTENT** **PAGE**

**Table S1. Enzymes cocktails used to produce structural motifs from WAX.**

**Table S1. Enzymes cocktails used to produce structural motifs from WAX.** The substrate (WAX; 0.5 mg/mL) diluted in an appropriate buffer was incubated with the carbohydrate-active enzymes (CAZy) and hydrolysates were collected at 0.5 and 24 h after incubation with each enzyme.

| Cocktail | Substrate | Enzyme(s) | Buffer [pH/temperature, °C] | Collection points |
|---|---|---|---|---|
| 1 | WAX | AFASE | NaOAc [4.0/40 °C] | 2 |
| 2 | WAX | XYLNP | $Na_3PO_4$ [6.0/50 °C] | 2 |
| 3 | WAX | AFASE + XYLNP | NaOAc [4.0/40 °C] + $Na_3PO_4$ [6.0/50 °C] | 4 |
| 4 | WAX | FAERU + AFASE | $Na_3PO_4$ [7.0/40 °C] + NaOAc [4.0/40 °C] | 4 |
| 5 | WAX | FAERU + XYLNP | $Na_3PO_4$ [7.0/40 °C] + $Na_3PO_4$ [6.0/50 °C] | 4 |

WAX: arabinoxylan from wheat flour (MW 56 KDa, Ara:Xyl = 38:62); AFASE: α-L-arabinofuranosidase purified from *Aspergillus niger*; XYLNP: endo-1,4-β-xylanase recombinant from *Neocallimastix patriciarum*; FAERU: feruloyl esterase (recombinant enzyme from rumen microorganism); NaOAc: 100 mM sodium acetate buffer; $Na_3PO_4$: 100 mM sodium phosphate buffer.

**Resource. OneLab platform - SPE-PGC purification of WAX structural motifs**

The ".onp" file available in the supplementary material can be upload in the "OneLab design & execute" online platform (https://onelab.andrewalliance.com/login) and be used as a semi-automated protocol to purify oligomers through solid-phase extraction using porous graphitic carbon resin (SPE-PGC). A pipetting robot (Andrew+, Waters Corporation) equipped with a set of Sartorius Picus2 electronic pipettes (Helsinki, Finland) and the following labware was used:

- GlycoWorks™ HILIC µElution plate (Waters Corporation)

- 1.5 mL short thread vial with wide opening (VWR International, Stockholm) in a 48x vial holder (Waters Corporation)

- 96-well PS F-bottom clear microplate (Greiner Bio-One, Kremsmünster, Austria)

- 4 x 10 mL multichannel reservoir (Integra Biosciences, Brøndby, Denmark)

**Fig S1. Changes in pH and short-chain fatty acids (SCFA) throughout *in vitro* colonic fermentation.**

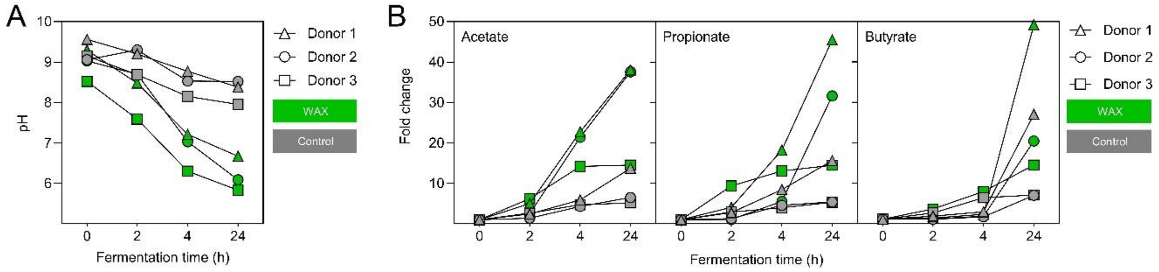

**Fig S1. Changes in pH and short-chain fatty acids (SCFA) throughout *in vitro* colonic fermentation.** (**A**) pH and (**B**) relative levels of the SCFA acetate, propionate and butyrate throughout *in vitro* colonic fermentation of *Kruskakli* (source of wheat arabinoxylan, WAX). Control: *in vitro* colonic fermentation without substrate. Samples were collected at time 0 (10 min), 2, 4 and 24 h. Fermentation was performed in three batches using the faecal inoculum of different donors.

### Table S2. Limit of quantification (LOQ) of oligomers in samples obtained from *in vitro* fermentation.

**Table S2. Limit of quantification (LOQ) of oligomers in samples obtained from *in vitro* fermentation.** The source of arabinoxylan (WAX) was subjected to *in vitro* colonic fermentation at a concentration of 10 mg/mL, resulting in a concentration of 200 µg/mL WAX. A 500 µL-aliquot was further fractionated, concentrated and quantified using the phenol-sulfuric assay method. The limit of quantification (LOQ) considers all concentration steps and is defined as the lowest value that can be measured with certainty using the phenol-sulfuric assay.

| Oligomers | Limit of quantification (LOQ) | |
|---|---|---|
| | Fermentation extracts (µg/mL) | % (w/w) in relation to WAX |
| $A^3X$ | 0.11 | > 0.01 |
| $A^2XX$ | 0.02 | > 0.01 |
| $XA^3XX$ | 0.02 | > 0.01 |
| $XA^{2,3}XX$ | 10.43 | 0.52 |
| XTE | 26.13 | 1.31 |
| XPE | 27.76 | 1.39 |

**Table S3. Quantification of WAX in process blanks during human colonic fermentation in process blanks.**

**Table S3. Quantification of WAX in process blanks during human colonic fermentation in process blanks.** Structural motifs were quantified after purification, annotation, fractionation and concentration. Results are expressed as µg/ 100 mL of the oligomer in the fermentation extract (i.e., total amount of oligomer/fermentation flask). No WAX substrate was used in the process blank. LOQ: value below the limit of quantification (**Table S2**); –: not detected.

| **Oligomer fractions** | **Donor 1** | | | | **Donor 2** | | | | **Donor 3** | | | |
|---|---|---|---|---|---|---|---|---|---|---|---|---|
| | **0 h** | **2 h** | **4 h** | **24 h** | **0 h** | **2 h** | **4 h** | **24 h** | **0 h** | **2 h** | **4 h** | **24 h** |
| $A^3X$ | – | – | – | – | LOQ | – | – | – | – | – | – | – |
| $A^2XX$ | – | – | – | – | – | – | – | – | 16.1 | – | – | – |
| $XA^3XX$ | LOQ | LOQ | – | – | – | – | – | – | 27.8 | – | – | – |
| $A^{2,3}XX$ | 53.4 | LOQ | – | – | – | – | – | – | – | – | – | – |
| XTE | – | – | – | – | – | – | – | – | – | – | – | – |
| XPE | – | – | – | – | – | – | – | – | – | – | – | – |
| *Total* | 53.4 | – | – | – | – | – | – | – | 43.9 | – | – | – |

**Fig S2. Molecular docking of structural motifs from WAX with TLR2 and TLR4.**

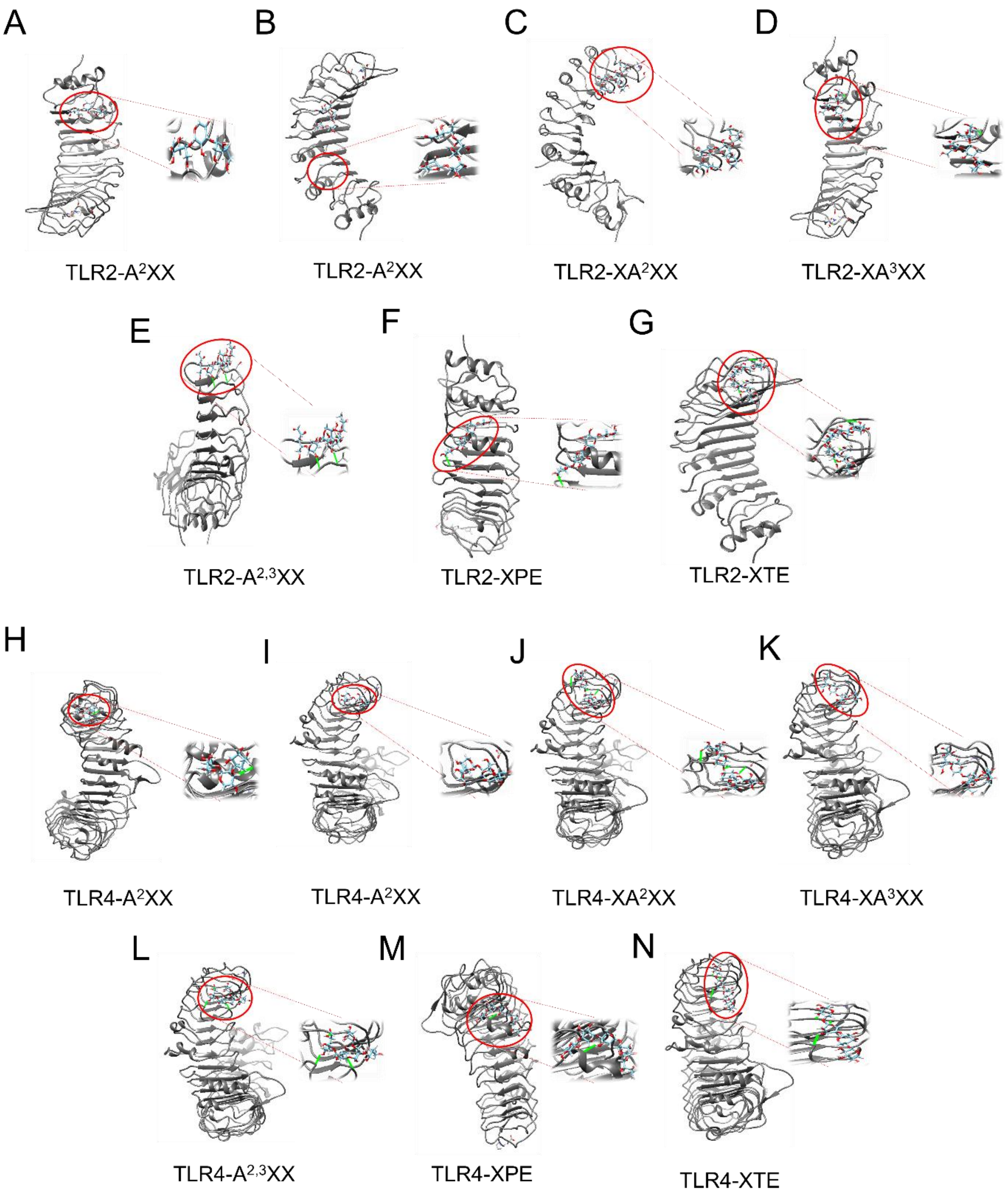

**Fig S2. Molecular docking of structural motifs from WAX with TLR2 and TLR4. Representative image of the highest score position of A$^{3}$X, A$^{2}$XX, XA$^{2}$XX, XA$^{3}$XX, A$^{2,3}$XX, XPE and XTE in the molecular docking model with (A-G) TLR2 and (H-N) TLR4.** XA$^{2}$XX and XA$^{3}$XX belong to the same structural motif fraction but were analysed separately to explore whether differences in the spatial conformation of structural motifs impact their binding capacity.
